# Nirmatrelvir Resistance in SARS-CoV-2 Omicron_BA.1 and WA1 Replicons and Escape Strategies

**DOI:** 10.1101/2022.12.31.522389

**Authors:** Shuiyun Lan, Grace Neilsen, Ryan L. Slack, William A. Cantara, Andres Emanuelli Castaner, Zachary C. Lorson, Nicole Lulkin, Huanchun Zhang, Jasper Lee, Maria E. Cilento, Philip R. Tedbury, Stefan G. Sarafianos

## Abstract

The antiviral component of Paxlovid, nirmatrelvir (NIR), forms a covalent bond with Cys145 of SARS-CoV-2 nsp5. To explore NIR resistance we designed mutations to impair binding of NIR over substrate. Using 12 Omicron (BA.1) and WA.1 SARS-CoV-2 replicons, cell-based complementation and enzymatic assays, we showed that in both strains, E166V imparted high NIR resistance (∼55-fold), with major decrease in WA1 replicon fitness (∼20-fold), but not BA.1 (∼2-fold). WA1 replicon fitness was restored by L50F. These differences may contribute to a potentially lower barrier to resistance in Omicron than WA1. E166V is rare in untreated patients, albeit more prevalent in paxlovid-treated EPIC-HR clinical trial patients. Importantly, NIR-resistant replicons with E166V or E166V/L50F remained susceptible to a) the flexible GC376, and b) PF-00835231, which forms additional interactions. Molecular dynamics simulations show steric clashes between the rigid and bulky NIR t-butyl and β-branched V166 distancing the NIR warhead from its Cys145 target. In contrast, GC376, through “wiggling and jiggling” accommodates V166 and still covalently binds Cys145. PF-00835231 uses its strategically positioned methoxy-indole to form a β-sheet and overcome E166V. Drug design based on strategic flexibility and main chain-targeting may help develop second-generation nsp5-targeting antivirals efficient against NIR-resistant viruses.

---

In 2019, SARS-CoV-2 emerged and began a rapid global spread, leading to the COVID-19 pandemic. As of December 2022, COVID-19 has caused 665 million cases and up to 6.7 million deaths worldwide [1]. This pandemic has also seen the advent of mRNA-based vaccines, which were rapidly produced and distributed, providing protection to millions of people [2-4]. Vaccines are, however, predominantly effective as preventative measures and do not typically help people who are already infected. Additionally, as the virus mutates and novel variants emerge [5, 6], vaccine efficacy may decline [7, 8]. Consequently, effective clinical control of the pandemic requires antivirals to treat infection and complement current prevention measures.

The first FDA-approved direct inhibitor of SARS-CoV-2 replication was remdesivir (RDV), which targets the viral RNA-dependent RNA polymerase or non-structural protein (nsp) 12 [9]. RDV was initially available in an injectable format, limiting its use to hospital settings; more recently an orally available prodrug of RDV has been reported [10, 11]. Two additional drugs have been approved and can be used in oral formulations: molnupiravir [12] and Paxlovid [13]. Molnupiravir inhibits SARS-CoV-2 replication through viral RNA mutation buildup [12]. A recent clinical trial reported that molnupiravir treatment does not significantly lower risk of hospital admission [14]. Paxlovid targets the protease nsp5, also known as the main protease (M^pro^) or 3-chymotrypsin-like protease (3CL^pro^). Paxlovid comprises nirmatrelvir (NIR) (Figure 1a), a tri-peptide-based antiviral that inhibits nsp5, and ritonavir, which improves the NIR pharmacokinetic profile. Nsp5 is responsible for cleaving 11 sites releasing nsp4 through nsp16 from polyproteins of nsps and contains a catalytic dyad Cys145-His41 that bears some similarities chymotrypsin [13]. Nsp5-targeting antivirals include GC376 and PF-00835231 (Figure 1a). GC376 is a dipeptide-based broad-spectrum inhibitor of coronavirus nsp5 proteases originally discovered as an inhibitor of feline coronaviruses shown to also inhibit SARS-CoV-2 nsp5 [15-18]. PF-00835231 is a ketone-based nsp5 inhibitor with potency and structure similar to GC376, albeit it is capped by a methoxy indole. While efflux transporter P-glycoprotein diminishes the efficacy of PF-00835231 in Vero E6 cells, it does not negatively impact its efficacy in either A549+ACE2 cells or human polarized airway epithelial cultures [19].

**Figure 1.**
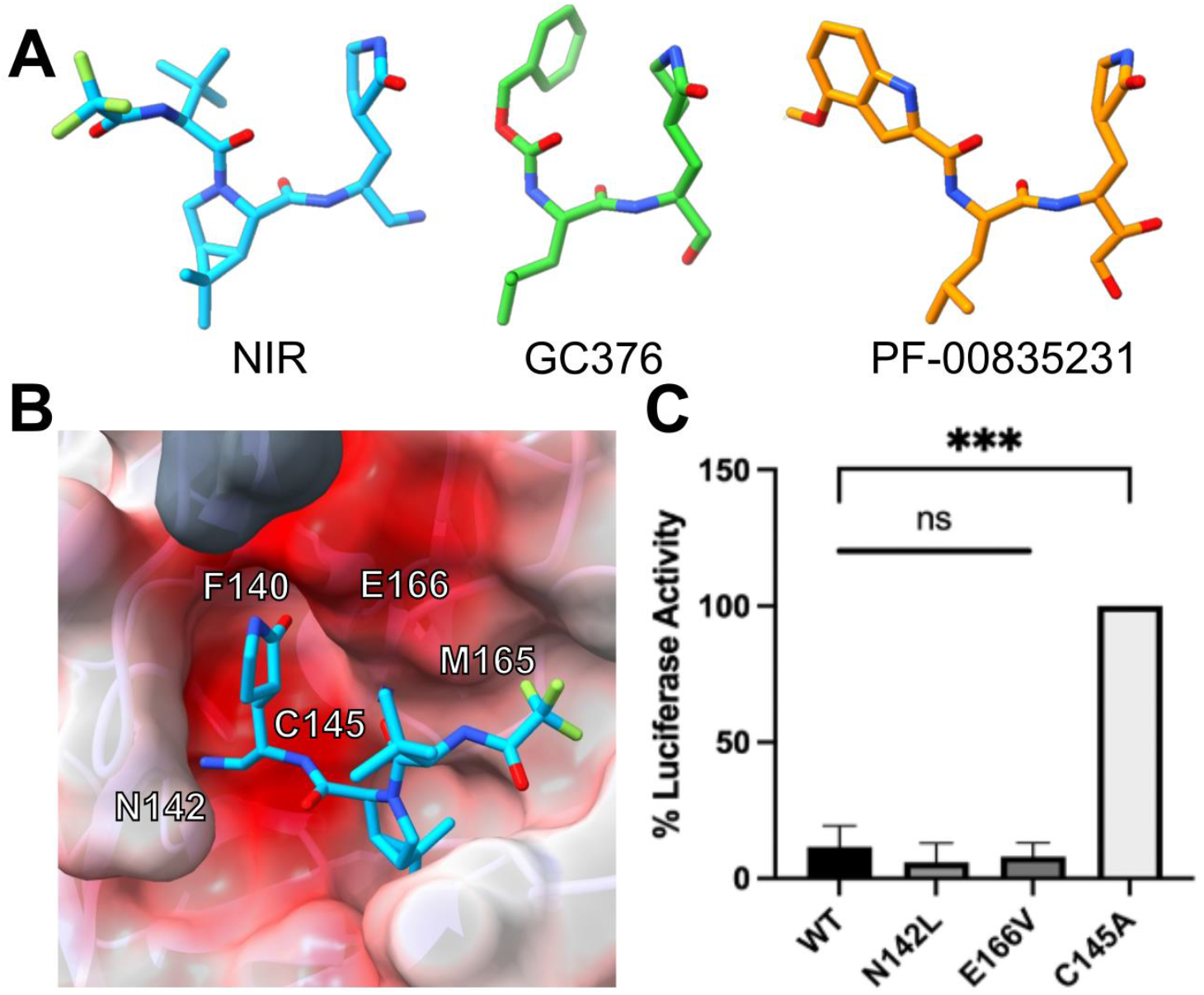
Preliminary analysis on impact of nsp5 mutations on protease activity and drug sensitivity. (a) Structures of nirmatrelvir (NIR), GC376, and PF-00835231 extracted from PDB ID: 7RFW, 7TGR, and 6XHM, respectively (without the respective warheads). (b) Close up of the nsp5 active site in complex with NIR (PDB ID: 7RFW). Color indicates electrostatics, and the dark grey indicates the N terminus of the other subunit. Residues F140, N142, M165, E166, and C145 were chosen for site directed mutagenesis. (c) Fitness of active site mutants measured in the absence of inhibitor. NanoLuc activity was normalized to the number of GFP (+) cells to control for transfection efficiency and the signal from the C145A nsp5. Both N142L and E166V showed lower NanoLuc activity (i.e., higher enzymatic activity) than the C145A nsp5 and comparable activity to the WT nsp5 (One-way ANOVA with Tukey p = 0.037 (n=2)).

NIR, GC376, and PF-00835231 mimic nsp5 substrates and covalently bind at the nsp5 active site through interactions between Cys145 and the inhibitor “warheads”: the cyano of NIR or the aldehyde of GC376, which is derived under physiological conditions from the bisulfite group [13, 15, 20], or the hydroxymethylketone of PF-00835231. Since receiving FDA approval, Paxlovid has been used to treat patients with COVID-19. There are several reports of SARS-CoV-2 rebound in Paxlovid-treated patients [21-25]. So far, reports of rebound have not been associated with resistance mutations. However, resistance could emerge if mutations at or near the active site of nsp5 impair NIR binding while still permitting cleavage of the natural substrates. Indeed, virological passaging data with NIR have already led to the investigation of various NIR-resistant mutations [26-29][59]. In this report, we test several possible mutations derived either from analysis of the structures of nsp5 bound to NIR or natural substrates [30, 31], or from reported data derived from passage of SARS-CoV-2 in the presence of NIR [26, 27][28]. We describe the effects of these mutations on fitness in replicon systems of Omicron and WA1 SARS-CoV-2 strains and test for drug resistance to NIR, GC376, and PF-00835231 [19, 32]. We also used molecular dynamics (MD) simulations to examine how different mutations may alter drug-protein interactions. Our results provide insight into potential mechanisms of NIR resistance and reveal structural attributes that can help design antivirals able to inhibit NIR (Paxlovid)-resistant SARS-CoV-2.

## Materials and Methods

### Cells

HEK293T/17 cells (#CRL-11268, American Type Culture Collection (ATCC), Manassas, VA, USA) were cultured in Dulbecco’s modified Eagle’s medium (DMEM, #10313-021, Gibco, Waltham, MA, USA) supplemented with 10% Serum Plus II (Sigma Aldrich, St. Louis, MO, USA), 100 U/ml penicillin, 100 μg/ml streptomycin (#400-109, Gemini Bioproducts, West Sacramento, CA, USA), and 2 mM L-glutamine (#25030-081, Gibco), in a humidified incubator at 37°C with 5% carbon dioxide.

### Plasmids

A previously described cell-based luciferase complementation reporter assay was used for initial assessment of SARS-CoV-2 nsp5 mutations and susceptibility to inhibitors [33]. We used versions of the nsp5-S-L-GFP reporter plasmid that were either wild-type (WT, of Washington or WA1 strain), catalytically inactive (C145A) or carrying mutations at other nsp5 positions (F140I, M165D, E166L, N142L, E166V, and E166I in the WA1 background). Mutants were generated by site directed mutagenesis using QuikChange II (Agilent, Santa Clara, CA, USA) and validated *via* Sanger sequencing (GeneWiz, Chelmsford, MA, USA).

SARS-CoV-2 replicons (SARS-2R_mNG_NeoR_NL) of WA1 and Omicron BA.1 were constructed either in the WT background (WT_WA1_, WT_Om_BA.1_) or in the presence of putative drug-resistance mutations at the 50, 132, 142, 166, and 167 sites of nsp5 as previously described [34]. All sequences were validated by long-read sequencing (Plasmidsaurus, Eugene, OR, USA). Construction of the N expression vectors was as previously described [34].

For biochemical assays, SARS-CoV-2 nsp5 was cloned into the pGEX-6P-1 vector using BamHI and XhoI and then synthesized commercially (GenScript, Piscataway, NJ, USA). A native N-terminus is attained during expression through an nsp5 autoprocessing site corresponding to the cleavage between nsp4 and nsp5 in the viral polyprotein, SAVLQ ↓ SGFRK, where ↓ denotes the cleavage site. The E166V mutation was introduced using QuikChange site-directed mutagenesis (Agilent, Santa Clara, CA, USA). Sequences were validated *via* Sanger sequencing (Azenta, Chelmsford, MA, USA).

### Cell-based luciferase complementation reporter assay

The nsp5-S-L-GFP reporter system contains the nsp5 sequence followed by a porcine teschovirus 2A cleavage signal and a NanoLuc luciferase sequence separated by the nsp4/nsp5 cut site [33]. GFP is also included to act as a transfection control. Functional nsp5 cleaves the nsp4/nsp5 site, rendering NanoLuc inactive; by contrast, inactive mutants do not cleave the nsp4/nsp5 site and NanoLuc activity can be measured. HEK293T/17 cells were seeded onto a 6-well plate and then transfected 24 h later using X-tremeGENE HP (Roche, Basel, Switzerland). After transfection (24 h) cells were re-seeded into a 96-well plate (40,000 cells/well) containing serial dilutions of inhibitors. After 24 h transfection efficiency was determined by counting GFP-positive cells. Cells were then lysed and NanoLuc activity measured using the NanoGlo Luciferase Assay System (Promega, Madison, WI, USA). Luciferase activity was normalized to the number of GFP positive cells in each well.

### Replicon fitness and dose response

SARS-CoV-2 replicons (SARS-2R_mNG_NeoR_NL) of WA1 and Omicron BA.1 strains were constructed in the absence of mutations (WT_WA1_, WT_Om_BA.1_) or in the presence of mutations at the 50, 132, 142, 166, and 167 sites of nsp5 as we have previously described [34]. HEK293T/17 cells seeded in a 6-well plate were transfected with 1 µg replicon plasmid (SARS-2R) using jetPRIME transfection reagent (Polyplus transfection, Illkirch-Graffenstaden, France). At 16 h post transfection, cells were trypsinized then seeded into 96-well plates and treated with serial dilutions of antivirals. NanoLuc luciferase assays were performed 48 h post treatment of SARS-2R with individual antivirals. Replicon fitness was determined by comparison of reporter gene expression by mutants to that of WT, when equal amounts of transfected nucleic acid were used (based on Nanodrop measurements). Dose response curves were calculated with Prism software (Graphpad, San Diego, CA, USA). All construct sequences were confirmed by Sanger sequencing (Azenta, Chelmsford, MA, USA) or full-length sequencing (Plasmidsaurus, Eugene, OR, USA). Sequencing results were analyzed with Lasergene/DNASTAR software (Madison, WI, USA).

### Expression and purification of nsp5 for biochemical assays

E166V_WA1_ was generated from the WT plasmid using QuikChange site directed mutagenesis. WT_WA1_ and E166V_WA1_ proteins were expressed in *E. coli* BL21 (DE3) by growing 50 mL LB cultures (containing 34 μg/mL chloramphenicol) to an OD_600_ of ∼0.8, induced by 0.2 mM isopropyl-thio-β-galactoside (IPTG) and incubated at 18℃ overnight with shaking. Bacteria were then pelleted by centrifuging for 30 min at 3,000 rpm, and pellets were kept at - 20℃ until purification.

Purification was performed using TALON resin. Bacteria pellets were resuspended in 10 mL of lysis buffer [25 mM Tris (pH 8.0), 300 mM NaCl, 5 mM β-mercaptoethanol (β-Me), and 4 mM MgCl_2_ with 1 mM phenylmethylsulfonyl fluoride (PMSF) and 0.15 mg/mL lysozyme] for half an hour then lysed using sonication. Cell debris were then pelleted by centrifugation (14,000 rpm at 4℃ for 30 min), and the supernatant was treated with 0.05% PEI before being centrifuged again. An equal volume of saturated ammonium sulfate was added to the supernatant and incubated at 4℃ overnight. Protein was pelleted by centrifugation (14,000 rpm at 4℃ for 30 min) and then resuspended in lysis buffer and centrifuged again before being added to 1-2 mL TALON resin (pre-equilibrated with lysis buffer). Supernatant and TALON resin were incubated at 4℃ for 2 h before being loaded onto a gravity flow column. The resin was then washed using lysis buffer with 0 mM, 20 mM, 50 mM, and 100 mM imidazole (pH 8.0) and eluted using lysis buffer with 300 mM imidazole (pH 8.0). The protein was then dialyzed and stored in 25 mM Tris (pH 8.0), 300 mM NaCl, 5 mM β-Me, and 4 mM MgCl_2_.

### Protease activity assay

Nsp5 activity was determined measuring changes in Fluorescence Resonance Energy Transfer (FRET) on peptide substrates carrying MCA (4-methylcoumaryl-7-amide) and DNP (2,4-dinitrophenyl) labels as previously described [35]. Measurements were performed in 20 mM Bis-Tris (pH 7.0) in a well volume of 100 μL of a peptide substrate that includes the nsp5 cleavage site between nsp4 and nsp5 proteins (nsp4-5) (-AVLQ ↓ SGFR[K(DNP)]K-*NH*_2_ (Millipore Sigma, Milwaukee, WI, USA; >95%). For WT_WA1_, 200 nM was used with 20 μM of substrate. For E166V_WA1_, 2 μM of the less active enzyme was used for increased signal. Activity was measured for up to 120 min on a Cytation 3 plate reader using a monochromator (Ex: λ = 322 nm / Em: λ = 381 nm). To determine the half maximal inhibitory concentration (IC_50_), the slope of the data from the linear region (the first 15 min) was determined and normalized to the slope of the uninhibited enzyme. These values were then used to calculate IC_50_s using GraphPad Prism 9.2.0 (GraphPad, San Diego, CA, USA).

### Sequence analysis

To ensure accurate and rigorous accounting of each amino acid change, two independent methods were used to analyze the EpiCoV database, the most comprehensive database of SARS-CoV-2 sequences currently available, which is curated by GISAID (global initiative on sharing all influenza data) database [36]. The first method used in-house python scripts (included in Supplementary Materials) to analyze the curated “allprot” protein sequence alignment obtained on 10/20/2022 from GISAID (Table 4, column 2). The “allprot” alignment is prepared by taking all sequences that have been submitted to EpiCoV and aligning their nucleotide sequences to the hCoV-19/Wuhan/WIV04/2019 (EPI_ISL_402124) reference sequence. From this alignment, the coding sequences for each protein is extracted and translated. From this dataset, in-house python scripts were used to filter out sequences that contain ambiguous residues (those for which one or more nucleotides were unassigned) and those that contain insertions and deletions. The filtered sequences (13,313,267 total sequences) were then screened for instances of specific amino acid changes related to this work. In the second method, the Coronavirus3D server [37] was used to identify instances of amino acid changes (Table 4, column 3).

### Molecular dynamics simulations

Initial structural coordinates for nsp5 in complex with GC376 (PDB ID: 7TGR), PF-00835231(PDB ID: 6XHM), and NIR (PDB ID: 7RFW) were retrieved from the Protein Data Bank (http://www.rcsb.org/) [13, 38]. These structures were prepared for molecular dynamics (MD) simulations using the Maestro modeling environment within the Schrödinger Software suite (Schrödinger Release 2022-2: Schrödinger, LLC, New York, NY, 2021). Briefly, the protein preparation workflow was used to add hydrogens, assign disulfide bonds, remove co-crystalizing small molecules and ions, and fill in missing side chains [39]. Hydrogen bond (H-bond) assignments were optimized to resolve overlap; protonation states were assigned using PROPKA [40]. For the respective ligands, protonation and charge states were calculated at pH: 7.4 ± 2.0 and the initial state of the ligand was selected based on calculating the number of hydrogen bonds and the Epik penalty score (Schrödinger Release 2022-2: Schrödinger, LLC, New York, NY, 2021) [41, 42]. Finally, a restrained minimization was performed using the OPLS4 forcefield [43]. The prepared nsp5-inhibitor complexes were then solvated in a 12 Å x 12 Å x 12 Å box, using the TIP3P water model [44]. Counterions were added to neutralize the charge of the system, and additional Na^+^ and Cl^-^ ions were added to a final concentration of 150 mM. MD simulations were performed using the Desmond MD simulation package within the Schrödinger Software suite (Schrödinger Release 2022-2: Desmond Molecular Dynamics System, D. E. Shaw Research, New York, NY, 2021. Maestro-Desmond Interoperability Tools, Schrödinger, New York, NY, 2021). The model systems were initially relaxed using Maestro’s default relax model system protocol and equilibrated with a 5 ns simulation run under isothermal–isobaric (NPT) ensemble conditions (temperature: 310 K, pressure: 1.01325 bar). The coordinates of these model systems were then used as the starting point for 100 ns runs. All simulations were performed with a 2 fs time step, and coordinates recorded at an interval of 20 ps. Simulation event analysis and simulation interaction diagram tools within Maestro were used for trajectory analysis.

## Results

### Initial analysis of nsp5 mutations using a cell-based luciferase complementation reporter assay

Using the crystal structures of WA1 nsp5, we designed mutations at residues that are proximal to the inhibitor binding site (Figure 1b) that might affect NIR resistance (including F140I, N142L, M165D, E166V, and E166L). To obtain initial insight on the impact of these nsp5 mutations on protease activity we used a luciferase complementation reporter assay [33]. Loss of protease cleavage function in this system leaves luciferase intact and results in high NanoLuc activity. Hence, the active site mutant C145A, which is reported to be catalytically inactive [45] showed indeed the highest NanoLuc activity (Figure 1c). The WT nsp5 is the most active protease, resulting in the most efficient cleavage of NanoLuc and thus lowest NanoLuc activity (Figure 1c). The NanoLuc activities of N142L and E166V nsp5 constructs were between those of WT and C145A (Figure 1c). The F140L, M165D, and E166L mutations displayed NanoLuc activity that was even higher than C145A (p < 0.001, p = 0.0008, and p = 0.0004, respectively based on a one-way ANOVA with Tukey), suggesting that these mutations render nsp5 inactive. Since N142L and E166V did not eliminate nsp5 activity, we selected these for further studies using Omicron and WA1 replicon systems. For comparison purposes we later expanded the ongoing analysis of replicon mutations to include L50F, E166A, and L167F that were listed in preprint studies that appeared at the later stages of our study [26-27].

### Impact of putative resistance mutations on the drug susceptibility and fitness of SARS-CoV-2 replicons

To assess the effect of residue changes on NIR resistance we constructed mutant versions of SARS-CoV-2 replicons on the Washington (WA1) and Omicron (Om_BA.1) strain backgrounds. Similar to the corresponding viruses, the sequences of the WA1 and Om_BA.1 replicons differ at 13 positions throughout various nsp genes, only one of which is in the nsp5 region (P132 _WA1_ *vs*. H132_Om___BA.1_). Specifically, nsp3: K38R, S1265del, L1266I, A1892T; nsp4: T492I; nsp5: P132H; nsp6: L105del, S106del, G107del, I189V, L260F; nsp12: P323L; nsp14: I42V.

#### Effect on susceptibility to antivirals

We tested the susceptibility of 12 replicons (WA1, Table 1; Om_BA.1, Table 2) in the presence and absence of putative drug-resistance mutations to several antivirals. We determined the EC_50_ values of RDV, NIR, GC376, PF-00835231. RDV targets nsp12. Hence, as expected, all replicons (WT_WA1,_ WT_Om___BA.1_ and all mutants in Tables 1 and 2) had comparable EC_50_s for this drug. In contrast, the WA1 mutants displayed varying degrees of resistance to NIR, 2.5-fold for N142L_WA1_, 53-fold for E166V_WA1_, 18-fold for the (L50F/E166A/L167F)_WA1_ triple mutant, and 109-fold for the (L50F/E166V)_WA1_ double mutant. Interestingly, the GC376 nsp5 inhibitor displayed similar activity against all the mutants, with only (L50F/E166A/L167F)_WA1_ showing a small increase in EC_50_ (∼3-fold). PF-00835231 has a similar resistance profile as GC376, fully inhibiting E166V_WA1_ and (L50F/E166V)_WA1_ and exhibiting a ∼9-fold decrease in efficacy against (L50F/E166A/L167F)_WA1_ (Table 1). Similar patterns are seen in the Om_BA.1 replicon with L50F/E166V showing more resistance than E166V, triple, and N142L to NIR, and only the triple (containing E166A, rather than E166V) showed some rather modest resistance to GC376 and PF-00835231 (Table 2). Additionally, the WT_Om_BA.1_ proved slightly more resistant than WT_WA1_ to all antivirals.

**Table 1.** Susceptibility of WA1 (Washington strain) SARS-CoV-2 replicons to nsp5 inhibitors remdesivir (RDV), nirmatrelvir (NIR), GC376, and PF-00835231. Replicon-transfected 293T cells were incubated with serially diluted compounds in 96-well plates for 48 h. Replication was assessed by Nanoluc activity. EC_50_ and standard deviations are indicated. RDV is an nsp12-targeting antiviral serving as control. ND: not determined.

| WA1 Washington strain | EC<sub>50</sub> ± SD / μM<br>(Fold change from respective WT<sub>WA1</sub>) |  |  |  |
| --- | --- | --- | --- | --- |
| nsp5 variant | RDV | NIR | GC376 | PF-00835231 |
| WT<sub>WA1</sub> | 0.010 ± 0.003<br>(1) | 0.034 ± 0.009<br>(1) | 0.21 ± 0.03<br>(1) | 0.13 ± 0.057<br>(1) |
| E166V<sub>WA1</sub> | 0.011 ± 0.002<br>(1.1) | 1.8 ± 0.6<br>(53) | 0.19 ± 0.02<br>(0.9) | 0.12 ± 0.13<br>(0.96) |
| (L50F/E166V)<sub>WA1</sub> | 0.007 ± 0.003<br>(0.7) | 3.87 ± 1.05<br>(114) | 0.49 ± 0.13<br>(2.3) | 0.066 ± 0.033<br>(0.51) |
| (L50F/E166A/L167F)<sub>WA1</sub> | 0.011 ± 0.001<br>(1.1) | 0.61 ± 0.05<br>(18) | 0.71 ± 0.03<br>(3.4) | 1.2 ± 0.49<br>(9.2) |
| (L50F/E166A/L167F/P132H)<sub>WA1</sub> | ND | 0.92 ± 0.33<br>(27) | ND | ND |
| N142L<sub>WA1</sub> | 0.009 ± 0.001<br>(0.9) | 0.084 ± 0.01<br>(2.5) | 0.32 ± 0.03<br>(1.5) | 0.76 ± 0.68<br>(5.8) |

**Table 2.** Susceptibility of Omicron BA.1 SARS-CoV-2 replicons to nsp5 inhibitors RDV, NIR, GC376, and PF-00835231. Replicon-transfected 293T cells were incubated with serially diluted compounds in 96-well plates for 48 h. Replication was assessed by Nanoluc activity. EC_50_ and standard deviations are indicated. RDV is an nsp12-targeting antiviral serving as control. ND: not determined.

| Omicron BA.1 strain | EC<sub>50</sub> ± SD / μM<br>(Fold change from WT<sub>BA.1</sub>) |  |  |  |
| --- | --- | --- | --- | --- |
|  | RDV | NIR | GC376 | PF-00835231 |
| WT<sub>Om_BA.1</sub> | 0.014 ± 0.002<br>(1) | 0.11 ± 0.02<br>(1) | 0.70 ± 0.24<br>(1) | 0.35 ± 0.10<br>(1) |
| E166V<sub>Om_BA.1</sub> | 0.006 ± 0.001<br>(0.4) | 6.01 ± 1.55<br>(54) | 0.41 ± 0.20<br>(0.58) | 0.14 ± 0.076<br>(0.40) |
| (L50F/E166V)<sub>Om_BA.1</sub> | 0.006 ± 0.001<br>(0.4) | 8.65 ± 0.47<br>(79) | 0.62 ± 0.12<br>(0.89) | 0.13 ± 0.02<br>(0.37) |
| (L50F/E166A/L167F)<sub>Om_BA.1</sub> | 0.009 ± 0.004<br>(0.7) | 2.92 ± 0.50<br>(26) | 3.8 ± 2.0<br>(5.4) | 6.3 ± 0.71 (18) |
| (L50F/E166A/L167F/H132P)<sub>Om_BA.1</sub> | ND | 4.8 ± 2.7<br>(44) | ND | ND |
| N142L<sub>Om_BA.1</sub> | 0.010 ± 0.001<br>(0.7) | 0.11 ± 0.02<br>(1.0) | 0.46 ± 0.14<br>(0.65) | 0.33 ± 0.071<br>(0.95) |

#### Effect on fitness

To assess the replication fitness of the 12 replicons, we compared the reporter gene expression to that of the corresponding WT in the absence of antivirals, when starting with equal amounts of nucleic acid replicons. N142L_WA1_ and E166V_WA1_ exhibited replication defects (∼5-fold and at least 10-fold, respectively) compared to WT_WA1_ (Fig. 2). Similar to others [27][28] we found that introducing L50F into the E166V replicon (L50F/E166V)_WA1_ restores fitness to wild-type levels in WT_WA1_ (Fig. 2). We found the triple mutant (L50F/E166A/L167F)_WA1_ (by Jochmans and colleagues [26]) replicated efficiently compared to WT_WA1_ (Figure 2). However, in the Om_BA.1 backbone, the E166V mutation did not decrease the fitness as much as it did in WA1 (E166V_Om___BA.1_ had ∼50% less activity of WT_Om___BA.1_, compared to ∼95% loss of activity of E166V_WA1_ compared to WT_WA1_) (Figure 2). Also, the L50F mutation had no effect on Om_BA.1 [E166V_Om___BA.1_ and (L50F/E166V)_Om___BA.1_ had comparable activities; Figure 2b]. Notably, in contrast to the WA1 strain, the (L50F_E166A_L167F)_Om___BA.1_ triple mutant appeared significantly decreased fitness compared to WT_Om___BA.1_. To determine whether the effect of the triple mutation on fitness was the result of the single mutation in nsp5 residue 132, we introduced mutations at 132 at the triple mutant background. Thus, we found that H132P in [(L50F/E166A/L167F/H132P)_Om___BA.1_ rescued the poor fitness of the triple mutant [(L50F/E166A/L167F)_Om___BA.1_. The effect was mirrored by introducing P132H in WA1 [(L50F/E166A/L167F/P132H)_WA1_], which suppressed the strong fitness of [(L50F/E166A/L167F)_WA1_] (Figure 2).

**Figure 2.**
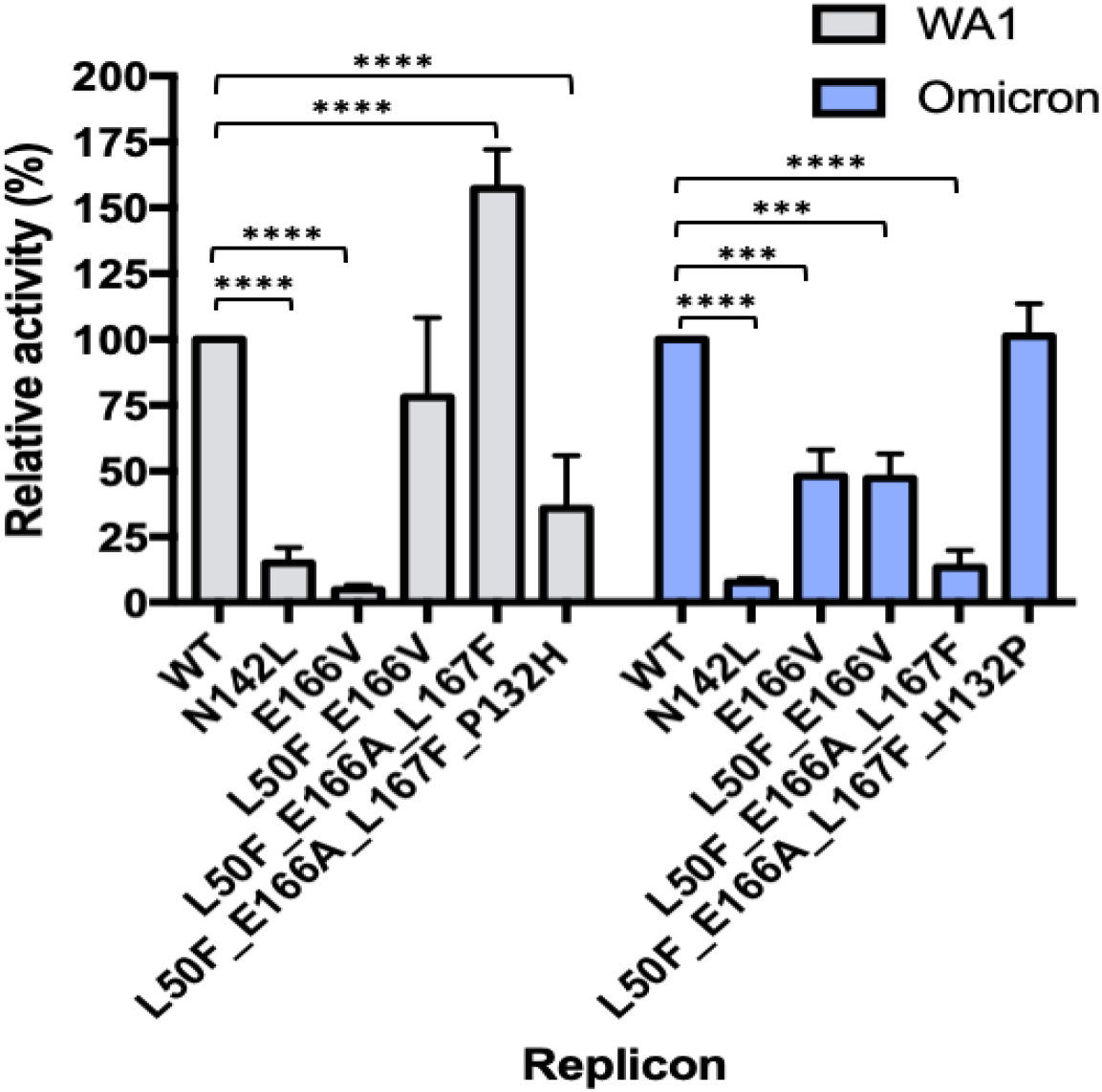
Impact of nsp5 mutations on replicon fitness of SARS-CoV-2 replicon mutants. 293T cells seeded in 24-well plate were transfected with 0.25 μg replicon plasmids together with 0.05 μg N expression vector. Nanoluc activities of WA1 (a) and Om_BA.1 (b) wild-type and mutant replicon strains were measured at 48 hpt. P values determined by 2-way ANOVA; ***, p < 0.001; ****, p < 0.0001.

### Resistance to nsp5 inhibitors in enzymatic protease activity assay

IC_50_ values derived from an *in vitro* assay reflect this same pattern of E166V-nsp5_WA1_ displaying significant resistance to NIR and decreased fitness compared to WT-nsp5_WA1_. An i*n vitro* activity assay was used to determine IC_50_ values for the WT-nsp5_WA1_ and E166V-nsp5_WA1_ proteins with both GC376 and NIR (Table 3). This assay uses a fluorescently labeled peptide of the nsp4-5 cleavage site to measure nsp5 activity. In this assay, the E166V-nsp5_WA1_ mutant showed significantly decreased activity, so a 10-fold higher concentration was used to increase the signal. Under these experimental conditions, the cleavage activity of E166V-nsp5_WA1_ resembled that of the WT-nsp5_WA1_ and The E166V-nsp5_WA1_ resistance to NIR was 23-fold higher than that of GC376.

**Table 3.** IC_50_ values of WT and E166V nsp5 based on the protease activity assay. Values represent average and standard deviations from n = 3 replicates.

| WA.1 nsp5 enzyme | IC<sub>50</sub>, μM<br>(Fold Change from WT<sub>WA1</sub>) |  |
| --- | --- | --- |
|  | NIR | GC376 |
| WT-nsp5<sub>WA1</sub> | 0.030 ± 0.009<br>(1) | 0.040 ± 0.02<br>(1) |
| E166V-nsp5<sub>WA1</sub> | 14.2 ± 6<br>(473) | 0.838 ± 0.2<br>(21) |

### Analysis of viral sequences submitted in the GISAID

Examination of the SARS-CoV-2 sequences (all strains included; October, 2022) through a search of these individual mutations in the GISAID database showed that all these mutations have very low prevalence, most likely as a result of their impaired fitness. However, L50F (and T21I [27]) occurred at a considerably higher prevalence (Table 4 and [27-28].

**Table 4.** Number of instances where each amino acid change has been reported in the GISAID database. Values are instances reported out of 13,313,267 sequences analyzed using either an in-house python script or the Coronavirus3D webserver.

| Amino Acid Change | Instances<sup>a</sup> | Instances<sup>b</sup> |
| --- | --- | --- |
| L50F | 4,986 | 4,013 |
| N142L | 17 | 5 |
| C145X | 37 | 7 |
| E166A | 9 | 6 |
| E166V | 10 | 0 |
| L167F | 7 | 0 |
| E166A/L50F | 1 | 0 |

<sup>a</sup> Instances found using in-house python scripts. <sup>b</sup> Instances found using the Coronavirus3D webserver [37]

### Molecular Dynamics Analysis

To better understand the effects of the nsp5 mutations on resistance to antivirals we conducted molecular dynamics (MD) simulations of multiple nsp5-inhibitor complexes. The canonical active form of nsp5 is a dimer [46], which at physiological conditions (pH ∼7.4) has each of the substrate binding sites positioned proximal to the N-terminal domain of the neighboring subunit. We initially ran pilot MD simulations of either the individual nsp5 monomers or of nsp5 dimers and did not observe significant structural differences between the active sites. Hence, we proceeded to study the following complexes in the dimer form, using simulations that were at least 100 ns in duration: WT-nsp5_WA1_:NIR, WT-nsp5_Om_BA.1_:NIR, WT-nsp5_WA1_:GC376, WT-nsp5_Om_BA.1_:GC376, E166V-nsp5_WA1_:NIR, E166V-nsp5_Om_BA.1_:NIR, E166V-nsp5_WA1_:GC376, E166V-nsp5_Om_BA.1_:GC376, WT-nsp5_WA1_:PF-00835231, E166V-nsp5_WA1_:PF-00835231. These simulations addressed the following questions:

#### E166V confers resistance to NIR

During the course of the simulations of the WT-nsp5_WA1_:NIR and E166V-nsp5_WA1_:NIR complexes, the inhibitors remained bound to the proteins and were generally well constrained at the active sites. NIR sampled a slightly narrower range of conformations in the WT rather than in the E166V mutant complexes (Figure 3a): specifically, the average changes in displacement of non-hydrogen NIR atoms had RMSD values of 1.2 Å *vs*. 2.3 Å for the WT-nsp5_WA1_:NIR and E166V-nsp5_WA1_:NIR complexes. In addition, inspection of the individual NIR atom positions (Figure 3b) revealed rather small changes in individual atom positions (typically ∼1 Å RMSF). Apparent exceptions include the RMSF peaks at (i) carbon atoms #16, #17, #18 in the tertiary butyl (t-butyl) group; (ii) fluorine atoms #35, #36, and #37 in the trifluoromethyl group (Figure 3c). However, these changes reflect the sampling of identical rotamers that result from free rotation between the #21-#23 and #5-#22 chemical bonds (red and green circular arrows in Figure 3c); as such, these conformations are equivalent and equally present in both the WT-nsp5_WA1_ and E166V-nsp5_WA1_ simulations (Figure 3b,c). (iii) We also observe smaller changes at atoms near the C145-H41 catalytic center, namely at the cyano group (#24, #2) and at the atoms of the lactam ring (#29, #1, #9, #6, #7, #25, #9) (Figure 3b,c). Overall, NIR moves largely as a rigid body during the simulations (Figure 3d,e), albeit with a limited local torsional change (orange circular arrow) in the case of E166V. This torsional change, however, results in a significant change in the interatomic distances of the catalytic C145_Sγ_ in relation to the reactive cyano carbon [C2 in (C)] during the simulations of the WT-nsp5_WA1_ *vs*. the E166V-nsp5_WA1_ NIR complex (from 3.5 Å to 6.2 Å, respectively; Figure 3d,e,f). Essentially identical results were obtained for the simulations with the nsp5_Om___BA.1_ enzymes that only differ from nsp5_WA.1_ by the P132H mutation.

**Figure 3.**
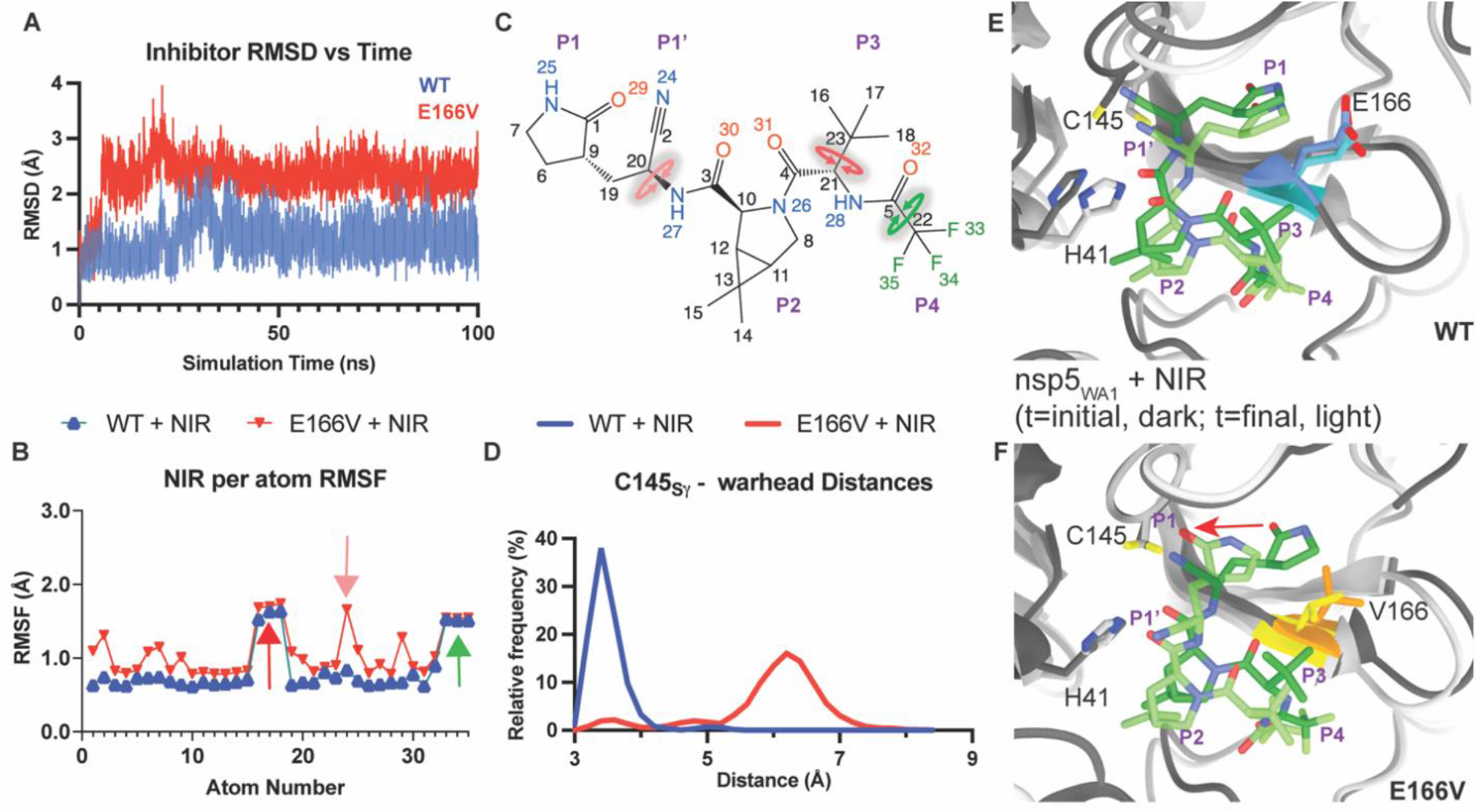
Conformational Rigidity Confers Changes in NIR Binding within the WT-nsp5_WA1_ and E166V-nsp5_WA1_ Active Sites. **(A)** Average Root Mean Squared Deviation (RMSD) of NIR binding in the WT- and E166V-nsp5_WA1_:NIR complexes, computed by aligning at every time point to the reference nsp5_WA1_ backbone structure at the beginning of the simulation, then computing the RMSD of non-hydrogen NIR atoms. The WT- and E166V-nsp5_WA1_:NIR simulations are in blue and red. **(B)** Positional Root Mean Squared Fluctuations (RMSF) of NIR atoms, numbered as in (C). Red and green arrows indicate the freely rotating P3 and P4 groups shown in (C). These values represent the internal atomic fluctuations of NIR at the end of the simulation compared to their starting positions. **(C)** 2D representation of NIR. Atom numbers correspond to the positions plotted on the horizontal axis in (B). Inhibitor sites are shown in purple, as previously defined. Rotating bonds corresponding to the colored arrows in (B) are indicated by circular arrows of the same color. **(D)** Frequency distribution (in percentage of total simulation time) of interatomic distances of the catalytic C145_Sγ_ in relation to the reactive cyano carbon [C2 in (C)]. **(E, F)** Superimposition of WT-nsp5_WA1_:NIR (E) and E166V-nsp5_WA1_:NIR (F) complexes at the beginning and end of the simulation (dark *vs*. light colors, respectively; E166 is in blue, V166 in yellow, rest of nsp5 in gray, NIR in green). Catalytic residues H41 and C145 are labeled. The red arrow indicates the lactam ring repositioning at the end of the simulation compared to its starting position.

#### GC376 evades resistance from the E166V mutation

The available crystal structures (6WTT, 8D4M, 8DD9, 8D4K [60]) and simulations of various nsp5:GC376 complexes suggest significantly different interactions at the inhibitor binding sites. GC376 is substantially more flexible than NIR (Figure 4a,b,c), primarily due to its P4 benzyl ester group, which can assume highly diverse conformations, with an RMSF for the positional variation of its aromatic ring atoms reaching 2.6 Å (blue arrow, Figure 4b), due to rotation of the P4 benzyl ester group around the bond between the C1 and O2 atoms (Figure 4c). Similarly, but to a lesser extent, the P3 Leu residue of GC376 can assume more conformations than the more constrained Leu analog P3 of NIR with the gem-dimethyl cyclopropane ring (RMSF Figure 4b,c). Hence, introduction of the inflexible V166 does not result in any significant repositioning of GC376 as a rigid body, because the inherent flexibility of GC376. Such flexibility is consistent with other experimentally determined structures [17, 47]. Consequently, the benzoate moiety at the P3 site helps defuse the potential steric conflict by almost freely repositioning within the active site during the simulations of WT-nsp5_WA1_:GC376 and E166V-nsp5_WA1_:GC376 (Figure 4b,c,e,f). Accordingly, there is no significant change in the interatomic distances of the C145_Sγ_ in relation to the aldehyde group warhead of GC376 during the simulations of the WT-nsp5_WA1_ *vs*. the E166V-nsp5_WA1_ NIR complex (Figure 3d,e,f). Essentially the same structural changes were observed in the simulations of the WT-nsp5_Om___BA1_ and E166V-nsp5_Om___BA1_ enzymes that only differ from the corresponding nsp5_WA.1_ enzymes by the P132H mutation.

**Figure 4.**
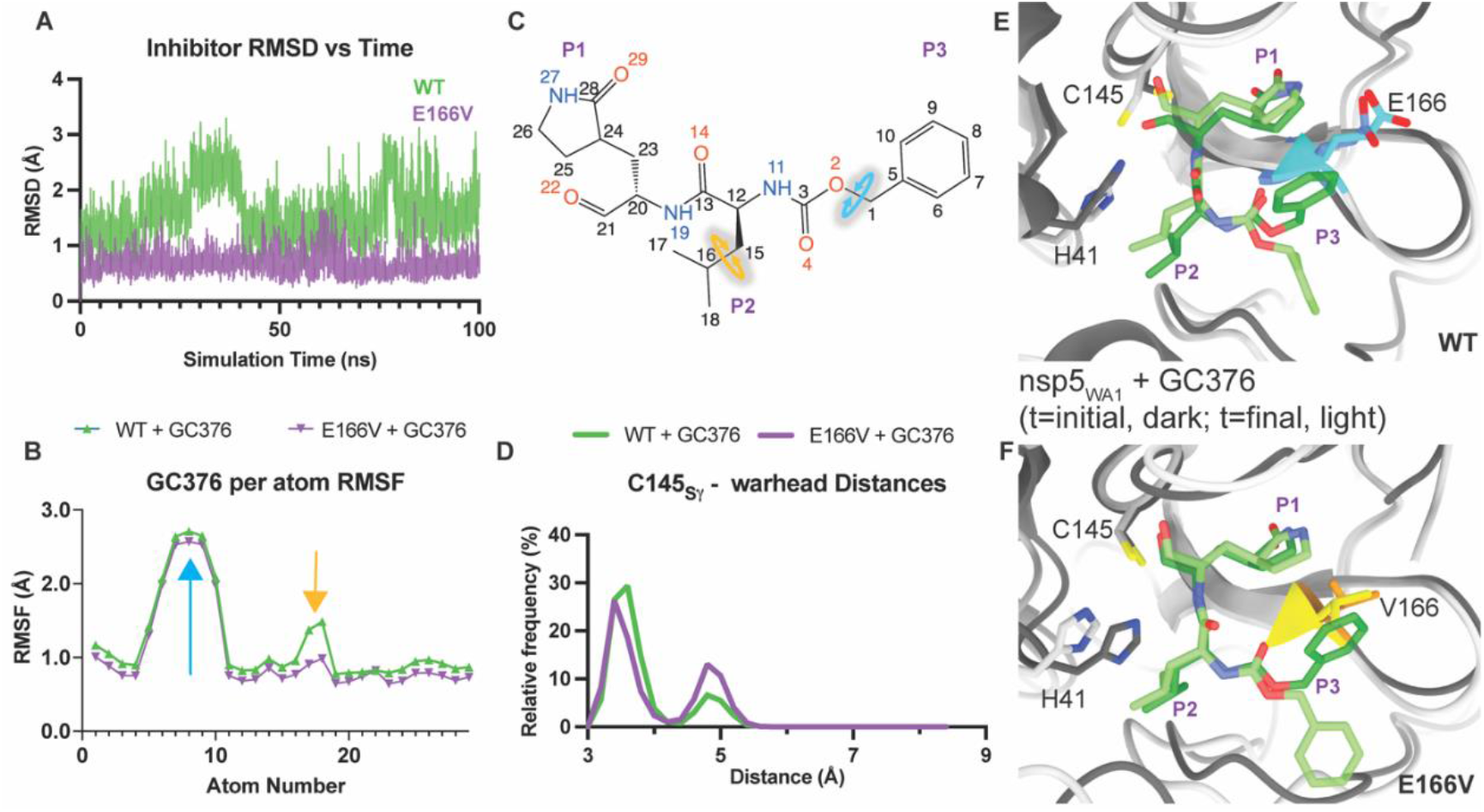
Structural Adaptation of GC376 within the WT-nsp5_WA1_ and E166V-nsp5_WA1_ Active Sites. **(A)** RMSD of GC376 binding in the WT- and E166V-nsp5_WA1_:GC376 complexes, computed by aligning at every time point to the reference nsp5_WA1_ backbone structure at the beginning of the simulation, then computing the RMSD of non-hydrogen GC376 atoms. The WT- and E166V-nsp5_WA1_:GC376 simulations are in green and purple. **(B)** Positional RMSF of GC376 atoms, numbered as in (C). Blue and yellow arrows indicate the relatively mobile P2 and P3 groups shown in (C). These values represent the internal atomic fluctuations of GC376 at the end of the simulation compared to their starting positions. **(C)** 2D representation of GC376. Atom numbers correspond to the positions plotted on the horizontal axis in (B). Inhibitor sites are in purple, as previously defined. Rotating bonds corresponding to the colored arrows in (B) are indicated by circular arrows of the same color. **(D)** Frequency distribution (in percentage of total simulation time) of interatomic distances of the catalytic C145_Sγ_ in relation to the reactive aldehyde carbon in GC376 [C21 in (C)]. **(E, F)** Superimposition of WT-nsp5_WA1_:GC376 (E) and E166V-nsp5_WA1_:GC376 (F) complexes at the beginning and end of the simulation (dark *vs*. light colors, respectively; E166 is in blue, V166 in yellow, rest of nsp5 in gray, NIR in green). Catalytic residues H41 and C145 are labeled.

#### PF-00835231 evades resistance from the E166V mutation

MD simulations showed that PF-00835231 remained bound to the proteins in the WT-nsp5_WA1_:PF-00835231 and E166V-nsp5_WA1_:PF-00835231 complexes (Figure 5a) without any major changes in individual atom positions during the course of the simulations (Figure 5b,c; RMSF variation ∼1 Å). As a result, there is no significant change in the interatomic distances between the C145_Sγ_ and the warhead of PF-00835231 and the targeted C145_Sɣ_ atom during the simulations of the WT-nsp5_WA1_ *vs*. the E166V-nsp5_WA1_ PF-00835231 complex (Figure 5d,e,f). Notably, the β-sheet-like interactions of the methoxy-indole group in the P3 site of PF-00835231 with the main chain peptide bond at residue 166 are maintained in the WT-nsp5_WA1_:PF-00835231 and E166V-nsp5_WA1_:PF-00835231 complexes during the MD simulations (Figure 5e,f), albeit with a slight twist of the ring methoxy-indole at the region near residues 166 and 167 of nsp5.

**Figure 5.**
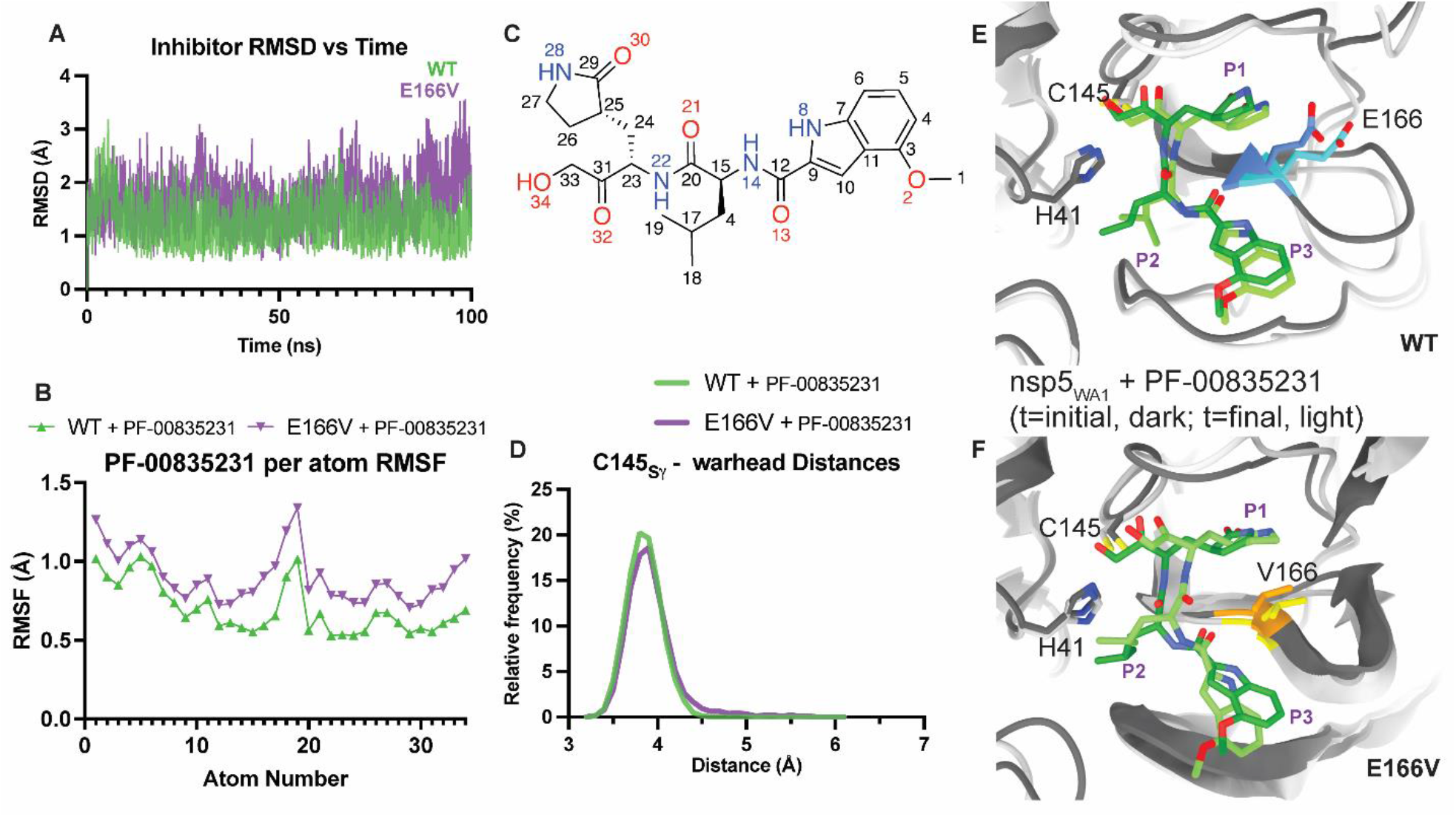
Conserved Binding of PF-00835231 within the WT-nsp5_WA1_ and E166V-nsp5_WA1_ Active Sites. **(A)** RMSD of PF-00835231 binding in the WT- and E166V-nsp5_WA1_:PF-00835231 complexes, computed by aligning at every time point to the reference nsp5_WA1_ backbone structure at the beginning of the simulation, then computing the RMSD of non-hydrogen PF-00835231 atoms. The WT-nsp5_WA1_:PF-00835231 and E166V-nsp5_WA1_:PF-00835231 simulations are shown in green and purple, respectively. **(B)** Positional RMSF of PF-00835231 atoms, numbered as in (C). **(C)** 2D representation of PF-00835231. Atom numbers correspond to the positions plotted on the horizontal axis in (B). Inhibitor sites are shown in purple, as previously defined. **(D)** Frequency distribution (in percentage of total simulation time) of interatomic distances of the catalytic C145_Sγ_ in relation to the reactive carbon in PF-00835231. **(E, F)** Superimposition of WT-nsp5_WA1_:PF-00835231 (E) and E166V-nsp5_WA1_:PF-00835231 (F) complexes at the beginning and end of the simulation (dark *vs*. light colors, respectively; E166 is in blue, V166 in yellow, rest of nsp5 in gray, PF-00835231 in green). Catalytic residues H41 and C145 are labeled.

#### *Effect of the difference in residue 132 between nsp5*_*Om_BA*.*1*_ *and nsp5*_*WA1*_

Overall, the P132H mutation observed in Omicron did not result in significant differences in the simulations of the nsp5 monomers of dimers or their substrate/inhibitor binding sites, consistent with the experimentally determined structures [31, 48]. Residue 132 is located in a loop between two β-strands, approximately 21-23 Å from the catalytic C145 for the duration of the simulations. Similarly, residue 132 is relatively distant from the nsp5 dimerization interface, only coming within 15-20 Å of the nearest residue (G283) in the neighboring subunit over the duration of the simulation. The only subtle local changes observed during the simulations of nsp5_Om_BA1_ compared to nsp5_WA1_ were at residues 115-117 that participate in β-sheet interactions slightly more frequently in simulations of the Om_BA.1 rather than the WT nsp5.

## Discussion

Due to the high selective pressure against the spike protein, new variants of the SARS-CoV-2 that escape current vaccines continue to emerge. Among the new strains, multiple versions of Omicron have been reported with more than 500 Omicron sublineages in existence [49-53][61]. Antiviral therapies are essential for treatment of those who are not vaccinated or cases of breakthrough infection. The nsp5 protease in SARS-CoV-2, and other coronaviruses, provides a promising drug target due to its essential role in coronavirus replication [49]. While Paxlovid (NIR-ritonavir) has shown promising efficacy [54] there are reports of viral RNA rebound in Paxlovid-treated COVID-19 patients [21-25] but so far, there is no direct link between post-treatment rebound and drug resistance mutations [29]. Nonetheless, the reported effects of nsp5 mutations on NIR resistance in cell culture experiments [29] suggest that drug-resistance mutations may become a threat to Paxlovid-based therapies, similar to antiviral treatments that target human immune deficiency virus, hepatitis B virus, influenza, and hepatitis C virus. This necessitates the design of second-generation inhibitors to combat viral strains resistant to NIR.

Regulatory constraints on gain-of-function studies on Paxlovid resistance development through *in vitro* passaging of infectious SARS-CoV-2, led us to alternative strategies for addressing the development and evasion of NIR resistance. We relied upon experience on the structural basis of antiviral resistance [55] and on efficient generation of relevant SARS-CoV-2 replicons [34] (>100, so far). We had previously demonstrated that rigid and bulky β-branched amino acid substitutions (such as M184V/I in HIV reverse transcriptase-RT) can sterically hinder binding of RT-targeting inhibitor over substrate, leading to viral resistance against FTC and 3TC, which are key drugs against HIV and HBV therapies [55]. We thus hypothesized that nsp5 active-site mutations could be identified that affect binding of NIR with limited effects on substrate binding, as it has been previously proposed by Schiffer and colleagues based on their protease inhibitor envelope hypothesis [56]. Indeed, using available structural data [30, 31] we tested several mutations and found E166V to impart strong NIR resistance initially using a cell-based luciferase complementation reporter assay (Figure 1c). We proceeded to construct twelve SARS-CoV-2 replicons, each containing one or more mutations. Unlike other NIR-resistance work focused on earlier variants (typically WA1), our constructed replicons belonged to the Omicron_BA.1 strains but also to WA1 for comparison purposes.

### Effects of mutations on Omicron_BA.1 and WA1 replicon fitness

Early on it became clear that the E166V mutation not only confers strong NIR resistance but also a significant loss of WA1 replicon fitness (∼95%), which can be restored to WT-nsp5_WA1_ levels by adding L50F (Figure 2). These findings are consistent with recent independently published work that used recombinant WA1 viruses [26][27][28] and also with comprehensive mutational analysis of nsp5 [62]. Surprisingly, while the Omicron_BA.1 replicons were also resistant to NIR (Tables 1, 2), we observed only a modest replication fitness loss (50%) for E166V-nsp5_BA.1_. While addition of a secondary mutation restored WA1 fitness in this and other studies (L50F, [27] and Figure 1, or T21I [28]), we did not observe change in the L50F/E166V-nsp5_BA.1_ fitness (Figure 2).

### Are Omicron_BA.1 data applicable to other Omicron lineages?

Currently, the vast majority of circulating viruses are of the Omicron strain [63], which as of 12/28/2022 includes BA.1, BA.3, BA.4, BA.5, BF.7, XBB, BQ.1, and their sublineages [63][64]. There are 13 residues that are different in the orf1ab sequences of BA.1 and WA1; there are also few additional non-structural protein residues that differ among various Omicron sublineages. However, our sequence analysis of Omicron genomes listed at the GSAID site show that their nsp5 sequences are more than 99% identical and have P132H mutation (https://gisaid.org). Given the differential effect of E166V on the fitness of WA1 *vs*. Omicron BA.1 replicons (Figure 2), we sought to determine the effect on NIR susceptibility and replicon fitness when we only mutate 1 of the 13 residues, leaving the other 12 unchanged. As an example, we examined the L50F/E166A/L167F replicon from the WA1 and Om_BA.1 strains: data in Figure 2 and Tables 1 and 2 showed that the observed phenotypic fitness differences between these replicons can be reversed with the single mutation at residue 132, without significantly affecting the resistance to NIR resistance. Specifically, addition of P132H to (L50F/E166A/L167F)_WA1_ resulted in a WA1 replicon with an Omicron-like nsp sequence [(L50F/E166A/L167F/P132H)_WA1_] that decreased its fitness (Figure 2). Conversely, the H132P change in the Omicron replicon (L50F/E166A/L167F)_BA.1_ converted its nsp5 to a WA1-like sequence (L50F/E166A/L167F/H132P)_BA.1_ and restored its fitness (Figure 2). Hence, these data suggest that the resistance and fitness data with the BA.1 Omicron replicon are likely to be applicable in other Omicron systems as well, which carry by overwhelming majority the phenotype-altering P132H mutation.

### Is the barrier to NIR resistance lower for Omicron than for WA1 strain viruses?

We hypothesize that the apparent strain-specific differences in replicon fitness of E166V NIR-resistant replicons may provide insight into the barrier to resistance among the studied strains. The poor fitness of E166V_WA1_ combined with the need for secondary mutations (L50F or T21I [26-28]) for sufficient viral growth is consistent with a relatively higher barrier to resistance compared to E166_BA.1_, which appears to have a significantly lower replicon fitness cost while maintaining NIR resistance. Consistent with the lower fitness of E166V-carrying viruses, this mutation has been reported only 10 times in 13,313,267 sequences submitted at the GISAID database (∼0.000075%). Interestingly, while peer-reviewed information on NIR resistance is not yet publicly available, Pfizer reported in the Paxlovid label that during the EPIC-HR clinical trial, E166V was found more often in NIR/ritonavir-rather than in placebo-treated individuals and that from the 361 drug treated patients, 3 had the E166V mutation (∼0.8%) [29]. Therefore, based on these early preliminary reports there seems to be a large (∼0.8% / ∼0.000075%, or ∼11,000-fold) increase in the probability of encountering the E166V mutation among treated patients compared to the total number of patients. To fully address this hypothesis, more detailed analysis of considerably larger future clinical studies will be required. Such studies would shed light on how often E166V appears among treated patients and how the type of variant may affect emergence of drug resistance.

### Molecular basis of differences in Omicron and WA1 fitness

The decreased fitness of E166V-nsp5 is likely due to the multiple functions of E166, whose interactions with substrates and inhibitors are observed in numerous crystal structures and are important for the enzymatic activity of nsp5. The E166 side chain is engaged in a H-bond network with water molecules and Q143, which is part of the catalytically important oxyanion hole. E166 directly interacts with the lactam ring of NIR (Figure 1b). It also forms a H-bond with S1 of the neighboring protomer (Figure 1b) and is thus involved in dimerization, which affects enzymatic function [57, 58]. All these H-bonds are lost by the E166V mutation. Indeed, recent MD studies suggested that V166 would destabilize nsp5 dimerization through disruption of dimer interactions S1-F140 and R4-E290 [27]. Also, biochemical experiments in a preprint sugest interference of A166 on nsp5 dimerization [26]. To elucidate why E166V has dissimilar deleterious effects when nsp5 has a P132 (WA1) or H132 (BA.1), we conducted MD simulations that revealed subtle changes at residues 115-117 of nsp5. Of note, residue Y118 in this region, directly interacts with the nsp13/14 peptide substrate when co-crystallized with nsp5 (PDB ID: 7TBT) [56], raising the possibility of SARS-CoV-2 strain-specific differences in interactions with substrates that may lead to effects on fitness.

### NIR resistance and how to overcome it

Virological data in Tables 1 and 2 demonstrated that the E166V mutation imparts strong resistance to NIR (∼55-fold) in both BA.1 and WA1 replicons. Using an expanded set of replicons it was also shown that resistance was even stronger in L50F/E166V (up to >100-fold). MD analysis indicated that NIR binds the active site as a relatively, yet not completely, rigid body during the simulations (Figure 3d,e). While this notable rigidity of NIR imparts strong binding and is consistent with the low nM antiviral EC_50_s (Tables 1 and 2), it comes, however, at the expense of conformational flexibility. Simulation of the E166V-nsp5_WA1_:NIR complex suggests sterically driven repositioning of the bulky NIR t-butyl (P3 group) by the also rigid and bulky beta-branched V166 (Figure 3d). While relocation of the P3 t-butyl alleviates the steric conflict with V166, the overall rigidity of NIR causes repositioning of the lactam ring (P1 group) at the catalytic site and concomitant reorientation of the NIR P1’ cyano group warhead. These changes appear to also affect the position of catalytic residue H41. Thus, the rigid-body movement of NIR, which is the result of its limited conformational flexibility, may also affect the efficiency of the interactions between the warhead and its C145 target, possibly affecting the NIR covalent binding at the E166V-nsp5_WA1_ active site. This negative impact may be further augmented by the loss of the H-bond interactions of E166, which deform part of the active site and also likely affect the affinity of NIR binding, as recently suggested [27]. Based on this model, we predict that other β-branched mutants (E166T, E166I) may also have similar effects on NIR resistance.

### GC376 evades E166V-based NIR-resistance through strategic torsional flexibility and structural adaptation or “wiggling and jiggling”

Virological data in Tables 1 and 2 show that unlike the strong NIR resistance (>50-fold to >100-fold) conferred by E166V or L50F/E166V in both BA.1 and WA1 replicons, there is a striking lack of resistance of the same replicons against GC376 (0.6-2-fold). MD simulations (Figures 3 and 4) show sharp differences in the flexibility of NIR *vs*. GC376. The flexible region of GC376 that significantly moves during the course of the simulation is the P4 benzyl ester group, strategically proximal to the bulky and inflexible V166, but also able to assume highly diverse conformations (Figure 4f). The conformational variability that helps GC376 avoid steric conflict with V166, is not only observed in MD simulations, but also in experimentally determined similar nsp5:GC376 structures (^17,47^ and PDB ID: 6WTT *vs*. 8D4M, 8DD9, 8D4K). Hence, strategic design of antivirals with flexible, adaptable structures could be a helpful approach to minimize steric hindrance-based drug resistance. A similar strategy known as “wiggling and jiggling” had been proposed by Arnold and colleagues in the design of non-nucleoside reverse transcriptase inhibitors (NNRTIs) that can bind in multiple conformations at the evolving NNRTI-binding pocket of HIV reverse transcriptase and avoid drug resistance mutations [66].

### PF-00835231 evades E166V-based NIR-resistance through strategic substitutions that enable interactions with conserved elements of the active site

Similar to GC376, PF-00835231 is essentially fully active against E166V or L50F/E166V in both BA.1 and WA1 replicons (Tables 1 and 2). Intriguingly, MD simulations showed that PF-00835231is not as flexible as GC376, and yet it is still able to maintain binding at E166V-containing active sites and inhibiting the corresponding viruses. We propose this is due to the β-sheet-like interactions of the methoxy-indole group in the P3 site of PF-00835231 with the main chain peptide bond at residue 166, which are maintained in the WT-nsp5_WA1_:PF-00835231 and E166V-nsp5_WA1_:PF-00835231 complexes during the MD simulations (Figure 5e,f). Such strategic placement of substitutions that introduce new stabilizing interactions may help overcome resistance. Some modest resistance to PF-00835231 is observed (∼9- to 18-fold) when 3 mutations are introduced at the inhibitor binding sites (L50F/E166A/L167F), likely due to interactions of the inflexible methoxy-indole with the bulky L167F, as proposed in a preprint [26]. Further increasing the flexibility of the methoxy indole in future analogs might help avoid additional steric clashes with L167F and help entirely suppress the mild resistance in the case of L50F/E166A/L167F. While the clinical development of the prodrug of PF-00835321 has been discontinued [65] there are valuable lessons learned for designing compounds that can avoid the key mechanisms of NIR resistance.

In conclusion, we independently identified and characterized a mutation that confers strong NIR-resistance in both Omicron and WA1 strains. We showed differences in the fitness of replicons, which may lead to variability in barriers to resistance between Omicron and non-Omicron strains. We report that two nsp5-targeting antivirals maintain potency against NIR-resistant replicons of Omicron and non-Omicron strains, and propose specific strategies for designing second generation antivirals against NIR-resistant viral strains.

## Acknowledgements

We acknowledge cloning support by the Emory Integrated Genomics Core (EIGC), which is subsidized by the Emory University School of Medicine and is one of the Emory Integrated Core Facilities. Additional support was provided by the National Center for Advancing Translational Sciences of the National Institutes of Health under Award Number UL1TR000454. The content is solely the responsibility of the authors and does not necessarily reflect the official views of the National Institutes of Health. SGS acknowledges funding support from NIH (R01-AI167356) and from the Nahmias‐Schinazi Chair fund at Emory University. We are grateful to Wei-Shau Hu and Vinay Pathak (HIV Dynamics and Replication Program, Center for Cancer Research, NCI at Frederick) for the nsp5-S-L-GFP reporter system. R.L.S. and G.N acknowledge funding support from the National Institute of Allergy and Infectious Diseases (T32-AI157855 and T32-GM008367, respectively). We thank Qinfang Sun, Abhishek Thakur, and Ronald M. Levy for helpful discussions and assistance in setting up Molecular Dynamics simulations and Karen A. Kirby for help editing the manuscript and useful discussions. We are grateful to Raymond F. Schinazi and Keivan Zandi, for the original stock of the WA1 strain and to Ann Chahroudi and Nils Schoof for the BA.1 strain stocks that were used to construct replicons. We also thank Eleftherios Michailidis for many useful discussions. Finally, we are grateful to Jindi Lan (deceased) for his steadfast support of Shuiyun Lan and family, enabling this research to continue during the COVID‐19 pandemic and shutdowns.

## Supplementary materials

Python script used to calculate frequency of different mutations in sequences in the EpiCoV database

~~~
# This code calculates the conservation of Nsp5 sequences in the “allprot”
file from GISAID.
# Uses the following criteria: must have 306 residues, no undefined
residues (“X”) or sequencing errors (“*”).
# Run program as follows: “python conservation.py [FastaFile]
[OutputFile]”
# Import required dependencies.
import os
import time
import sys
from Bio import SeqIO
import numpy as np
from numpy import genfromtxt
import pandas as pd
# Declaring command line variables
FastaFile = sys.argv[1] #
OutputFile = sys.argv[2]
# Defines a list of amino acids. resList = [“A”,”C”,”D”,”E”,”F”,”G”,”H”,”I”,”K”,”L”,”M”,”N”,”P”,”Q”,”R”,”S”,”T”,”V”,” W”,”Y”] # Defines the array that will be populated with conservation data ConArray = np.zeros((seqLen,20),dtype=int) # A basic counter for looping and the sequence length of Nsp5 counter = 0 seqLen = 306 # Function for processing a sequence into the conservation array def sequence_conservation(array):
~~~

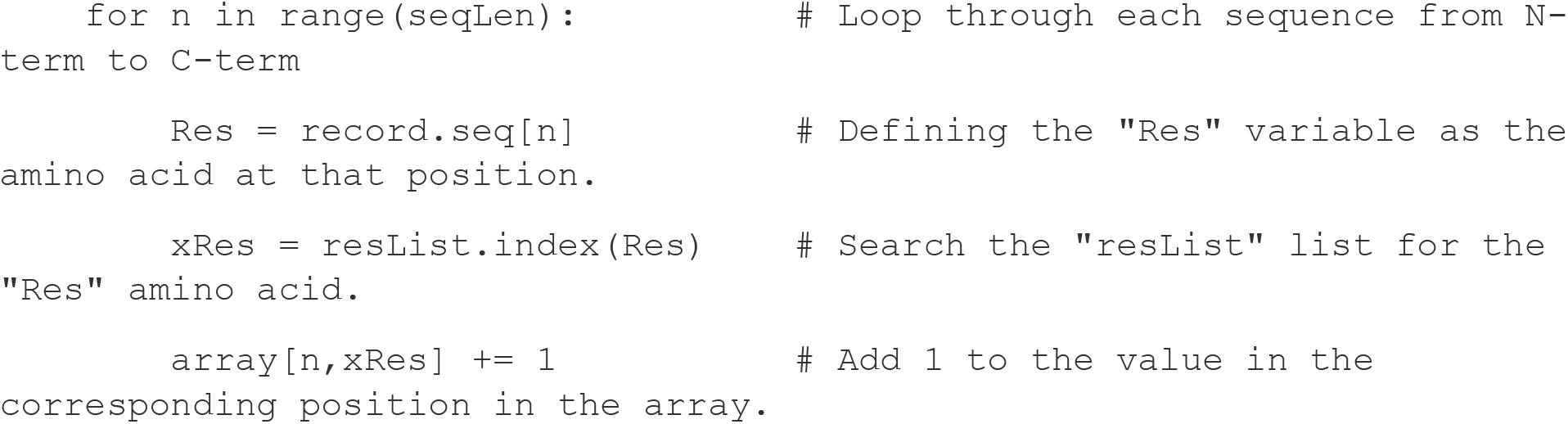

~~~
# Loop through the sequences in the fasta file.
for record in SeqIO.parse(FastaFile, “fasta”):
~~~

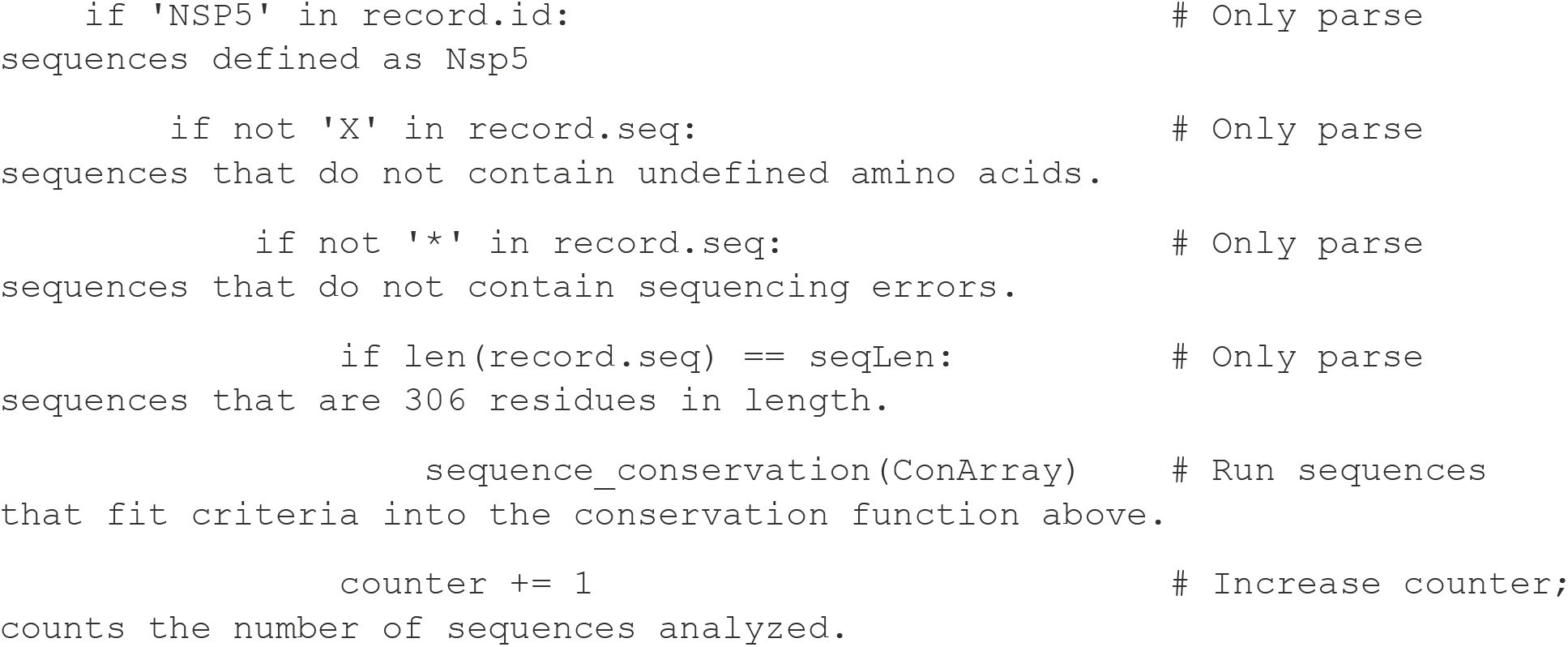

~~~
# Output array to a comma separated text file.
np.savetxt(OutputFile, np.array(ConArray), delimiter=‘,’, fmt=‘%s’)
# Print a completed statement to the command line.
print(f’Completed: [FastaFile] and saved array to [OutputFile]’)
~~~

